# Draft Genome Sequences of 10 Strains of *Pseudomonas syringae* pv. *actinidiae* Biovar 1: A Major Kiwifruit Bacterial Canker Pathogen in Japan

**DOI:** 10.1101/2020.06.25.170415

**Authors:** Takashi Fujikawa, Hiroe Hatomi, Hiroyuki Sawada

## Abstract

Several groups (biovars) of the kiwifruit bacterial canker pathogen, *Pseudomonas syringae* pv. *actinidiae*, are found in Japan. Here, we sequenced and compared the 10 genomes of biovar 1, the major group in Japan, which is known as the phaseolotoxin producer.

The kiwifruit bacterial canker pathogen, *Pseudomonas syringae* pv. *actinidiae* (Psa), was first described in Japan in 1989 (1). Subsequently, Psa was found in other kiwifruit-producing countries (2). Based on comparative analyses (2–4), Psa was categorized into several groups (biovars). The first Japanese group was named biovar 1 (Psa1), which was also later found in Italy and Korea. This biovar produces phaseolotoxin (2), a phytotoxin that inhibits arginine biosynthesis in host plants and results in bacterial canker symptom development. On the Psa1 chromosome, a large number of genes involved in phaseolotoxin biosynthesis are accumulated in an approximately 23 kb region (*argK-tox* cluster), which is contained in an exogenous genomic island (*tox* island) that Psa1 acquired in the past (2). However, some Psa1 strains found in Ehime Prefecture, Japan (the Ehime isolates) do not produce phaseolotoxin, although they seem to possess the *argK-tox* cluster (5). On the other hand, several Psa1 strains preserved in the NARO Genebank (https://www.gene.affrc.go.jp/index_en.php) may lack this cluster (2). Here, we selected 10 strains (Table 1) that represent Psa1 diversity and conducted comparative genome analyses.

---

Genomic DNA was extracted from the strains using the DNeasy mini kit (Qiagen, Hilden, Germany). Genomic DNA was sequenced using an Ion PGM sequencer with an Ion PGM Hi-Q View OT2 kit, an Ion PGM Hi-Q View Sequencing kit, and a 318 Chip kit v2 (all from Thermo Fisher Scientific Inc., Waltham, MA, USA). The sequence reads were quality controlled (a quality score < 20) and adapter sequences were removed using CLC Genomics Workbench version 12 (Qiagen). Using these reads, contigs (filtered with a size longer than 500 bp) were assembled *de novo* using the same software with default parameters (mapping mode = Create simple contig sequences (fast), automatic bubble size = yes, minimum contig length = 500, automatic word size = yeas, performing scaffolding = yes, auto-detect paired distances = yes). The draft genomes were annotated using the NCBI Prokaryotic Genome Annotation Pipeline (PGAP) v4. 1 (6).

The guanine and cytosine (G+C) content and genome size for these strains were found to be 58.2–58.8% and 4.9–6.3 Mbp, respectively (Table 1). PGAP identified 5,606–6,432 genes, including multiple rRNA and tRNA genes. In addition, the *argK-tox* cluster of each strain was sequenced (Table 2) and compared with the reference genome (CM002753) of ICMP 9617 (pathotype strain of Psa), indicating that some strains have synonymous substitution (silent mutation) in the cluster. Moreover, in the Ehime isolates (MAFF 211981 and 211983), it was clarified that a frameshift mutation in the fatty acid desaturase gene occurred due to a single G insertion, resulting in the loss of the ability to produce phaseolotoxin. On the chromosomes of MAFF 613017, 613018, and 212324, the *tox* island containing the *argK-tox* cluster could not be found, suggesting that these strains may not have experienced the island acquisition event. The fact that such diversification has occurred in the *argK-tox* cluster is an important piece of evidence for elucidating the pathogenicity, ecology and evolution of Psa1.

**Table 1.**
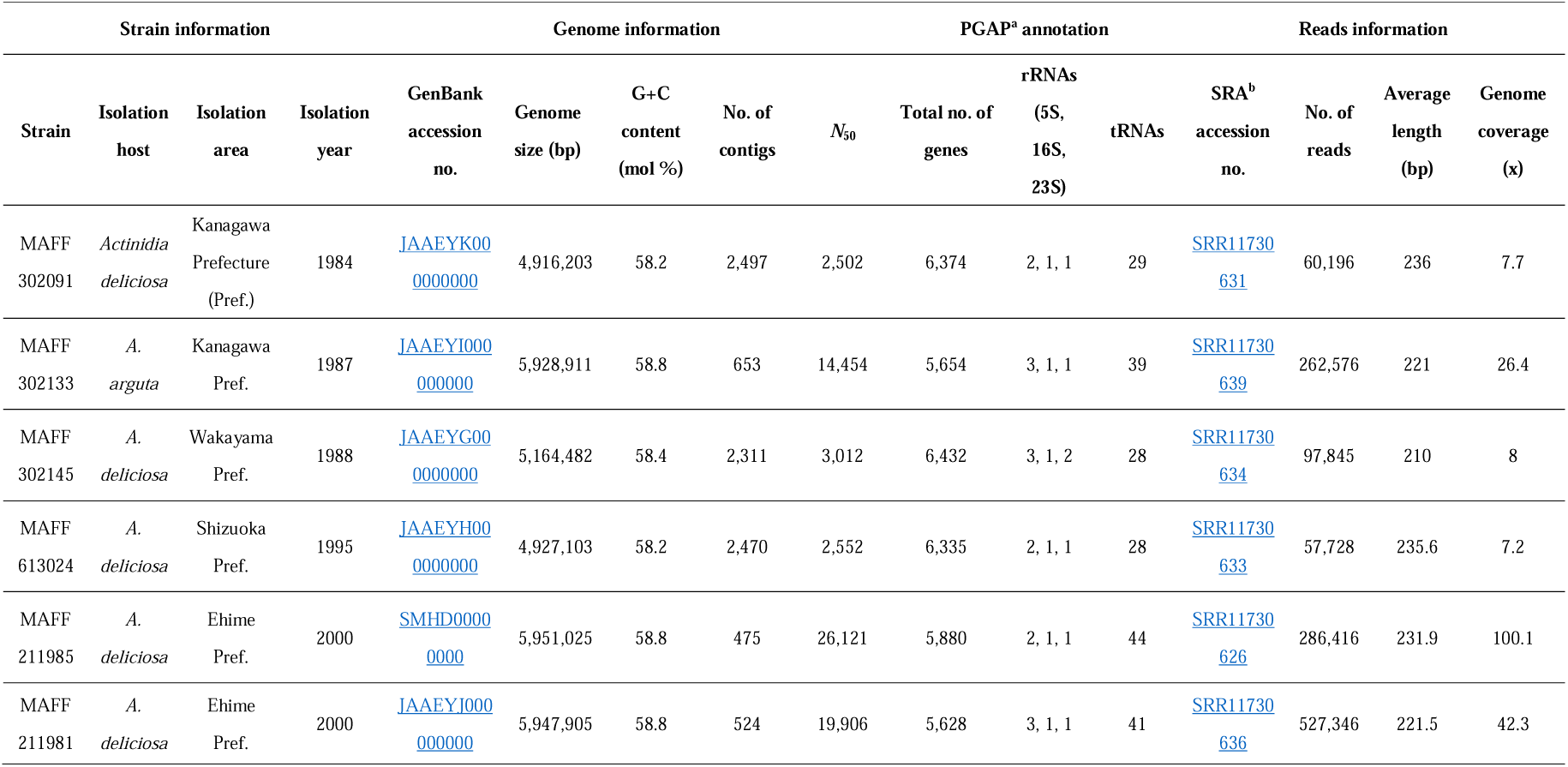
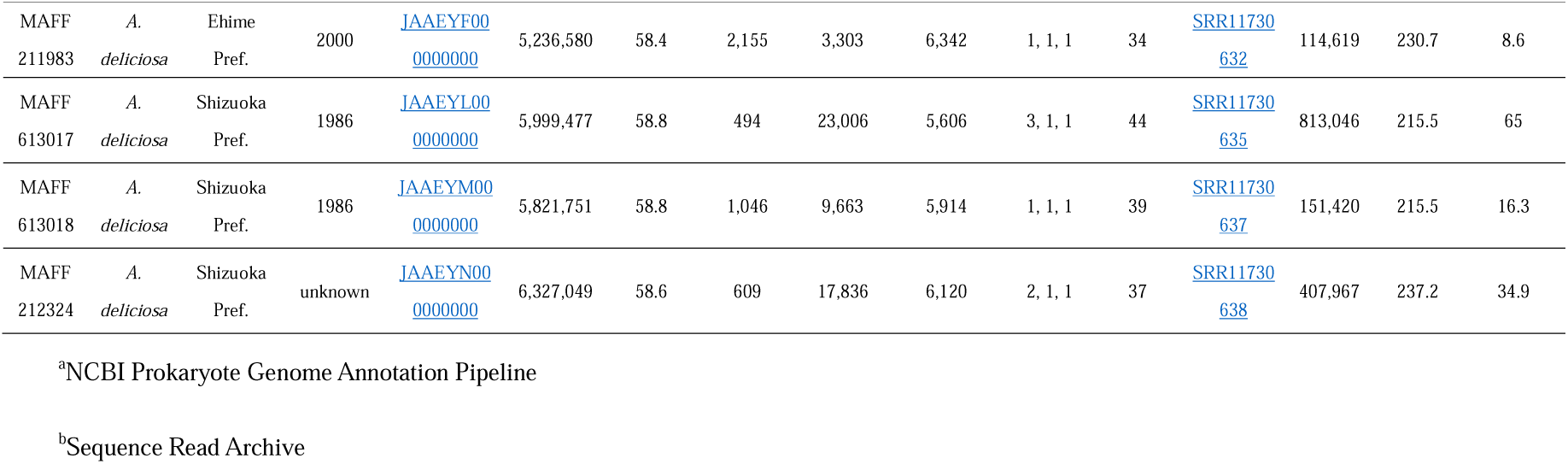
Genome data and accession numbers of 10 strains of *Pseudomonas syringae* pv. *actinidiae* biovar 1.

**Table 2.** Detail and accession numbers for the *argK-tox* gene cluster of 10 strains of *Pseudomonas syringae* pv. *actinidiae* biovar 1.

| *argK-tox* gene cluster information<sup>a</sup> |  |  |  |  |
| --- | --- | --- | --- | --- |
| Strain | Phaseolotoxin synthesis | *argK-tox* gene cluster | *argK-tox* gene cluster accession no. | Detail |
| MAFF 302091 | + | + | MT551019 | Same sequence as ICMP9617 |
| MAFF 302133 | + | + | MT551015 | Same sequence as ICMP9617 |
| MAFF 302145 | + | + | MT551014 | Same sequence as ICMP9617 |
| MAFF 613024 | + | + | MT551013 | Same sequence as ICMP9617 |
| MAFF 211985 | + | + | MT551017 | Synonymous substitution (silent mutation) in some coding genes against ICMP9617 |
| MAFF 211981 | - | + | MT551016 | Synonymous substitution (silent mutation) in some coding genes against ICMP9617<br>Frameshift mutation in the fatty acid desaturase gene due to the insertion of a single 'G' |
| MAFF 211983 | - | + | MT551018 | Synonymous substitution (silent mutation) in some coding genes against ICMP9617<br>Frameshift mutation in the fatty acid desaturase gene due to the insertion of a single 'G' |
| MAFF 613017 | - | - | - | Absence of *tox* island |
| MAFF 613018 | - | - | - | Absence of *tox* island |
| MAFF 212324 | - | - | - | Absence of *tox* island |

<sup>a</sup> + denotes presence, - denotes absence

## Data availability

All sequences identified in this study have been deposited in GenBank (see Tables 1 and 2 for accession numbers).

## Acknowledgments

We are grateful to Ms. M. Taguchi and Ms. A. Sasaki for supporting this work. We also thank the members of IFTS-NARO and GRC-NARO for their helpful discussions. We would like to thank Editage (www.editage.jp) for English language editing. This research received no specific grant from any funding agency in the public, commercial, or not-for-profit sectors.

## Notes

### Competing Interest Statement

The authors have declared no competing interest.

